# The transcription factor CLAMP is required for neurogenesis in *Drosophila melanogaster*

**DOI:** 10.1101/2020.10.09.333831

**Authors:** Maria A. Tsiarli, James Kentro, Ashley M. Conard, Lucy Xu, Erica Nguyen, Erica N. Larschan

**Affiliations:** Department of Molecular Biology, Cellular Biology and Biochemistry, Brown University, Providence, RI 02912, USA; Department of Neurobiology, Harvard Medical School, Boston, MA Department 02115, USA; Center for Computational and Molecular Biology, Brown University, Providence, RI, 02912; Computer Science Department, Brown University, Providence, RI, 02912

## Abstract

Neural stem cell (NSC) differentiation is controlled by cell-intrinsic and external signals from the stem cell niche including niche surface glia (SG). However, the mechanisms by which transcription factors drive NSC differentiation within the niche remain largely unknown. Here, we show that the *Drosophila melanogaster* transcription factor Chromatin-linked adaptor for MSL proteins (CLAMP) is required for regulation of stemness and proliferation of NSCs, especially in the optic lobe (OL). Using a validated *clamp* null mutant, we show that CLAMP promotes transcription of genes involved in stemness, proliferation, and glial development and represses transcription of genes involved in neurogenesis and niche survival. Consistent with transcriptional changes, CLAMP promotes NSC proliferation and niche SG production, while lack of CLAMP severely and specifically impacts OL development. Το identify potential mechanisms by which CLAMP may regulate brain development, we examined CLAMP binding site motifs and available CLAMP ChIP-seq data to determine which genes may be direct versus indirect targets. We found that genes encoding *Notch and tailless,* both master regulators of brain development, are directly bound by CLAMP, indicating that they are key direct targets of CLAMP. Moreover, *Notch* and *Tailless* are critical for OL development, providing a mechanistic link to the severe deficits observed in OL development in the *clamp* null mutants. Overall, our results suggest that CLAMP regulates a core transcriptional program which drives NSC proliferation and differentiation *via* cell-intrinsic and niche-dependent mechanisms involving niche glia and transcriptional regulation of *Notch* and *Tailless*.

## Introduction

Brain development is an extremely complex yet carefully orchestrated process, regulated along spatiotemporal axes. For the formation of functional brain circuitry to take place, neural stem cells (NSCs) must undergo multiple rounds of divisions and lineage decisions at each step of the process which are governed by specific transcription factors (TFs) (Konstantinides *et al*. 2018). In addition to the intrinsic developmental program, external metabolic and paracrine signals constitute another layer of developmental control over NSCs. Specifically, NSCs are part of a specialized microenvironment, the stem cell niche (Scadden 2006), which plays an architectural and nurturing role for the NSCs and their progeny. Moreover, the niche serves as a hub for the relay of metabolic and developmental signals that regulate differentiation of NSCs in order for the proper sequence of developmental events to be maintained (Speder and Brand 2018). In mammals, the stem cell niche is composed of various cell populations including neurons, glial cells, resident immune cells, the microglia, and blood vessels which along with endothelial cells and glial cells form the blood-brain-barrier (Dani *et al*. 2019). In *Drosophila*, the stem cell niche consists of NSCs and NSC progeny, including glial cells of various types that are vital to the maintenance of the homeostatic balance required for proper neurogenesis to take place (Hindle and Bainton 2014; Spéder and Brand 2014; Bailey *et al*. 2015; Kanai *et al*. 2018; Speder and Brand 2018; Ramon-Cañellas *et al*. 2019; Yuan *et al*. 2020). However, little is understood about the gene regulatory mechanisms that generate the stem cell niche.

In *Drosophila melanogaster* neurogenesis occurs at two distinct stages: the embryonic and post-embryonic stages (Hartenstein and Campos-Ortega 1984; Prokop and Technau 1991). The *Drosophila* central nervous system (CNS) is composed of the central brain (CB), the optic lobe (OL) and the ventral nerve cord (VNC). During the embryonic stage, CB neural stem cells (neuroblasts in *Drosophila*) proliferate to generate primary neurons until they become quiescent at the end of the embryonic stage. The optic lobe primordium which will form the OL, is generated by epithelial cells of the posterior procephalic region between embryonic stages 11-13 (Ramon-Cañellas *et al*. 2018). Between stages 12 and 13, the OL primordium invaginates and attaches at the ventrolateral surface of the brain and embryonic OL neuroblasts (EONs) are formed (Hakes *et al*. 2018). EONs proliferate briefly to generate glial and neuronal progeny and also enter quiescence (Hakes *et al*. 2018; Contreras *et al*. 2019b). During the early larval stage, concurrent with the initiation of larval feeding, metabolic signals impinge on the NSCs and cause exit from quiescence and initiation of post-embryonic neurogenesis. Sequential expression of transcription factors in proliferating NSCs, including of *Tailless* and *Notch*, regulate the temporal generation of diverse progeny types (Li *et al*. 2013; Bertet *et al*. 2014).

In *Drosophila,* distinct CNS glial types are identified based on morphological, positioning and molecular profile criteria: perineurial (PG), subperineurial glia (SPG), cortex glia (CG), astrocyte-like and ensheathing glia (Yildirim *et al*. 2019). The first two types of glia constitute the *Drosophila* brain-blood barrier and along with CG cells are also classified as surface glia due to their proximity to the brain surface (Contreras *et al*. 2019b). PG serve a nurturing role for the development and function of the NSCs and regulate NSC proliferation at different stages of larval development through intrinsic, metabolic and stem-cell niche mechanisms (Sousa-Nunes *et al*. 2011; Spéder and Brand 2014; Kanai *et al*. 2018) (Bailey *et al*. 2015; Kis *et al*. 2015). Importantly, CG form a bespoke membrane chamber around dividing NSCs and their progeny, thus providing protection to the proliferating NSCs and their niche (Speder and Brand 2018), whereas *mir8*^+^ surface-associated CG, control neurogenesis in the OL (Morante *et al*. 2013; Ramon-Cañellas *et al*. 2018; Yildirim *et al*. 2019). Thus, niche glia are important for NSC function. However, the molecular mechanisms that direct the functional and nurturing relationship between niche glia and NSCs remain poorly understood.

Here we define a role for the essential CLAMP zinc finger TF (Soruco *et al*. 2013b) in neurogenesis which is partly mediated by transcriptional regulation of key TFs controlling stemness and proliferation, and partly *via* niche glia. CLAMP (CG1832) was discovered as part of a screen for factors that are necessary for the function of the dosage compensation machinery in *Drosophila melanogaster* males (Larschan *et al*. 2012). CLAMP is essential for viability of both males and females and has transcriptional targets on all chromosomes, with a two-fold increased binding affinity on the X-chromosome compared to autosomes (Soruco *et al*. 2013b; Urban 2017). Since its first characterization as a required link for the dosage compensation complex to bind to the X-chromosome (Soruco *et al*. 2013a; Tikhonova *et al*. 2019), CLAMP is emerging as a multifaceted transcription factor with roles including chromatin organization (Kaye *et al*. 2017), histone locus function (Rieder *et al*. 2017) and regulation of Pol II release from pausing (Urban *et al*. 2017). The identification of *Clamp*, as a functional interactor of *embryonic lethal abnormal visual system* (*elav*) and *erect wings* (*ewg),* critical regulators of embryonic and post-embryonic neural development, suggested a potential neuronal role of CLAMP (Haussmann *et al*. 2008; Haussmann and Soller 2010b). Therefore, we hypothesized that CLAMP is a TF which regulates neuronal development and NSC biology.

To test our hypothesis, we investigated the role of CLAMP in nervous system development during the *Drosophila* larval stage using a CRISPR/Cas9-generated frameshift mutant in the *clamp* gene which eliminates the expression of the CLAMP protein (Urban 2017). We show that CLAMP is important both for neuronal and glial cell generation and regulates NSC proliferation *via* cell-intrinsic and niche-dependent mechanisms. Specifically, *clamp* null mutants have smaller brains than controls, with the most prominent defect observed in the optic lobe. To determine the transcriptional networks affected by CLAMP, we generated mRNA-sequencing data from *clamp* mutant embryos and larval brains, followed by gene ontology (GO) and binding motif analysis which revealed CLAMP motifs at key neurogenesis genes and the glial-development regulator, *glial cells missing.* Further solidifying CLAMP’s role in controlling NSC biology, embryo ChIP-seq data analysis revealed that CLAMP directly binds and regulates the master neurogenesis and NSC function regulators *Notch* and *Tailless*.

In *Drosophila*, the Notch pathway is crucial for NSC formation and differentiation, regulating the stem cell state by controlling quiescence and reactivation, and determining the type of neuronal progeny via temporal patterning. The transcription factor Tailless (Tll) is an orphan nuclear receptor crucial for early brain patterning, NSC maintenance, and optic lobe development. Especially in the OL, Notch and Tll interact to control NSC function and fate specification(Xu *et al*. 2024). Thus, to further explore the *in vivo* relationship between CLAMP and Tailless, we conducted immunostaining in *clamp* null mutant brains using NSC, glia markers and Tailless, revealing a brain-wide reduction in proliferating NSCs, surface glia, and, a profound defect in the development of the OL, similar to the phenotype seen in *Tailless* knock-down brains (Guillermin *et al*. 2015). Furthermore, knock-down of *clamp* in glia but not in neurons, revealed that NSC proliferation and glia generation are partly regulated *via* glial CLAMP. Our results support a model in which CLAMP regulates a transcriptional program which drives NSC proliferation and differentiation *via* cell-intrinsic and niche-dependent mechanisms that involve direct transcriptional regulation of *Notch* and *Tailless* and a function in niche glia.

## Materials and Methods

### Fly husbandry

Fly stocks used in this study are as follows: *y^1^w^1^*wild type controls, *clamp^2^* null, and *CLAMP* rescue lines, which were generated by Urban et al.(2017) and described therein *(Urban 2017)*. Briefly, the *clamp* null is a CRISPR-Cas9 generated frameshift mutation which abolishes CLAMP protein expression, whereas the *clamp* rescue was generated by introduction of a full CLAMP rescue construct on chromosome 3L. For the RNAi experiments the following lines were used: *repo-GAL4*, *elav^c155^* (*elav-GAL4*) and UAS-*clamp-RNAi,* all obtained from the Bloomington Stock Center. All crosses were done at 25°C and a 12-hour light/dark cycle.

*Clamp* is maternally deposited and its expression is high during the majority of embryogenesis(Rieder *et al*. 2017) (Fig.1a). While the persistence of maternal *clamp* was not specifically evaluated in late embryos, CLAMP protein deposited by *clamp* heterozygous mothers persists beyond the embryonic stage and is present in the histone locus body throughout the development of the larval salivary gland (Rieder *et al*. 2017). Thus, it is possible that a trace amount of maternally deposited *clamp* could still be present at 16-18 hour embryos and reduces the strength of the zygotic *clamp* nulls phenotype compared with embryos that are depleted of maternal CLAMP, which die early in embryogenesis soon after zygotic genome activation.

### Immunostaining

The following antibodies were used: Rabbit anti-CLAMP (1:200, Abcam custom made), mouse anti-Repo (1:50, Developmental Studies Hybridoma Bank (DSHB) 8D12), rabbit anti-DCP1 (1:100, Cell Signaling), rat anti-Deadpan (Dpn, 1:100, Abcam), rat anti-Elav (1:50, DSHB), mouse anti-Prospero (Pros, 1:30, DSHB), PCNA (PC10, Abcam), Nc82 (1:10, DSHB), rabbit anti-Tailless (Kindly provided by Dr James Sharpe, EMBL-Barcelona). Secondary antibodies used were Alexa Fluor Conjugated A405, A488, A568, A647 (1:1000, Invitrogen). A standard staining protocol was used.

### Quantification of marker-positive and superficial glia cells

To quantify superficial glia, 3-μm projections (z-projections of three 1-μm optical sections) of the dorsal surface of third instar larval brain hemispheres stained for Repo were used and quantified manually using Adobe Photoshop, as exemplified by Kanai et al. (2018). Six hemispheres were quantified for each genotype from at least 3 animals. The same approach was used to quantify marker positive cells i.e., Dpn, Pros and EdU. To ensure consistency between samples of control and the *clamp* mutant brains basic morphological features based on DAPI staining in combination with stack depth were monitored.

### Image acquisition and processing

Fluorescent images were acquired using an inverted Olympus FLUOVIEW FV1000 confocal laser scanning microscope or a Leica TCS SP2 AOBS DMIRE2 inverted scanning confocal microscope. Adobe Photoshop was used to adjust brightness and contrast in images and ImageJ (or Fiji (Schindelin et al., 2012)) was used to analyze images, while Adobe Illustrator was used to compile figures.

### EdU Incorporation

EdU incorporation in proliferative brain cells, was performed using Click-It Edu Alexa Fluor A488 (Thermo Fisher Scientific, MA, USA) in an adaptation of the method detailed inTastan and Liu (2015) (Tastan and Liu 2015). Briefly, samples were dissected in ice-cold Grace’s media and then transferred in Grace’s media containing 5μg/ml of EdU, for 2 hours at room temperature. Then, samples were washed in 0.1% PBST and fixed in 4% PFA in 0.1% PBST for 20 min at room temperature. Next, tissues were washed in 5% HINGS in 0.1 % PBST and then permeabilized in 0.5% PBST for 20 min at room temperature. After this step, the EdU Click It steps were followed according to the manufacturer’s instructions and then additional antibody staining was followed as described above.

### Larval Staging

Third-instar larvae crawling on the vial walls were selected at mid/late stage using the bromophenol blue method adjusted from Maroni and Stamey (1983) (Maroni and Stamey 1983), using 0.05% w/v bromophenol blue-containing fly food. To control for variations in brain size due to crowding, experiments were done under controlled density conditions. Specifically, a set number of mated males (n=4) and females (n=6) were allowed to mate for 5 hours and then removed from the vial. Additionally, the progeny number was evaluated to validate that we did not have large deviations between control and *clamp* mutant larvae numbers.

### RNA extraction and RNA sequencing

Mid/late third instar larvae were staged using the bromophenol blue method and larval brains were dissected in ice-cold PBS, collected in TRIzol until RNA extraction was performed using the phenol-chloroform method. Each biological replicate was generated by pooling at least 12 larval brains. mRNA-focused sequencing libraries were constructed from total RNA using the Illumina Truseq RNA Library kit v2 according to the manufacturer’s instructions. Paired-end 100 bp or 50 bp sequencing was performed on an Illumina HiSeq2500 sequencer in High-Output mode at the MGH NextGen Sequencing Core. Four biological samples were used per experimental condition. As the control group, we used the CLAMP rescue line in which the wild-type CLAMP gene was inserted into an attP site under the control of its endogenous promoter (Urban 2017).

### RNA sequencing data analysis and Differential Gene Expression Analysis

RNA-seq reads were processed using FastQC (Andrews 2010) with default parameters to check the quality of raw sequence data and filter out sequences flagged for poor quality, and aligned using HISAT2 (Kim *et al*. 2015). Then read counts per gene were generated via HTSeq (Anders *et al*. 2015) using the *Drosophila melanogaster* genome dm6 (BDGP6) genome as reference. Differential gene expression analysis was performed using DESeq2 (Love *et al*. 2014). *gProfiler* was used for GO analysis (Raudvere *et al*. 2019). Using TIMEOR (Conard *et al*. 2020), the temporally differentially expressed genes were displayed in interactive expression cluster-maps along the developmental time scale (see Supplementary files: *Embryo_heatmap* and *L3_brain_heatmap*), from which additional, GO analysis was performed by TIMEOR *via* clusterProfiler (Yu *et al*. 2012) to evaluate for possible gene networks.

### Clamp Binding motifs

We used FIMO (Grant *et al*. 2011) to identify individual CLAMP motif occurrences within **±** 1KB from the transcription start and end sites for all genes within each gene trajectory cluster. The position weight matrix used (Kaye *et al*. 2018) highlights the CLAMP motif independently of GAF binding potential. The specific command used is: *(fimo --oc $OUTPUT_DIR/ CLAMP_PWM.txt $geneList)*

### Average profiles

The average control-normalized read depth across a set of regions was plotted using deepTools computeMatrix and plotProfile. The read depth source are ChIP-seq data in the ENCODE database for CG1832 (Clamp) from mixed sex embryos (0-24hrs), part of the modERN project (https://www.encodeproject.org/experiments/ENCSR858HYS/). The regions were chosen as each set of differentially expressed genes (DEGs) from the *clamp* null versus *clamp* rescue RNA-seq experiments in embryo and third instar larvae. These were kept as all DEGs in an experiment and as separate images for up or down regulated genes in the absence of *clamp*. The gene regions were scaled to match positions of the transcription start sites (TSS) and transcription end sites (TES) with 3K base pairs shown on either side of the gene.

### Western Blotting

Larval brains were dissected from wandering third-instar larvae in ice-cold 0.7% saline and transferred in PBS + 0.3% Triton X-100 on ice. Liquid was carefully removed, then samples were frozen in liquid nitrogen and stored at -80°C until use. Approximately ten brains were collected per biological replicate. For protein extraction, 50 μL of ice-cold RIPA buffer (150 μL1M Tris-HCl pH 8, 12 μL 0.5M EDTA, 0.003 g sodium deoxycholate, 15 μL 20% SDS, 90 μL 5M NaCl, 30 μL 50x protease inhibitor, and 2700 μL water) was added to each sample. Samples were homogenized using an electric handheld homogenizer and disposable plastic pestle. After homogenization, the pestle was rinsed with an additional 50 μL of RIPA for a total volume of 100 μL per tube. Samples were rotated at 4°C for 1-2 hours. Protein concentration was quantified using the Qubit Protein Assay kit (Thermo Fisher Scientific).

### Statistical Analysis

Excel and GraphPad Prism were used for statistical analysis. Bar graphs were generated using the mean and standard error of the mean (SEM) for each sample. Statistical significance was determined using an unpaired Student’s t-test.

### Data Availability Statement & Correspondence

Data and materials generated in this study are available upon request from the corresponding authors. Transcriptomic data have been deposited in Gene Expression Omnibus (GEO) Database, under the accession number GSE148704.

## Results

### CLAMP is expressed throughout the larval nervous system and in proliferating neuroblasts

CLAMP is maternally deposited at both the protein and RNA levels and therefore *clamp* transcript and CLAMP protein levels are present at high levels during early embryogenesis (Rieder *et al*. 2017). Near the end of embryogenesis (16-18 hours), during tissue differentiation, and the cessation of the first wave of neurogenesis, *clamp* transcript levels show a reduction followed by an increase between 20-24 hours and then a decrease throughout the post-embryonic neurogenesis period (Fig. 1a). Tissue-specific expression data from the modENCODE Development RNA-Seq Collection (Graveley *et al*. 2011; Brown *et al*. 2014)/Flybase (Thurmond *et al*. 2019) show that *clamp* is highly expressed in the third instar larval central nervous system (Fig. 1b).

**Figure 1.**
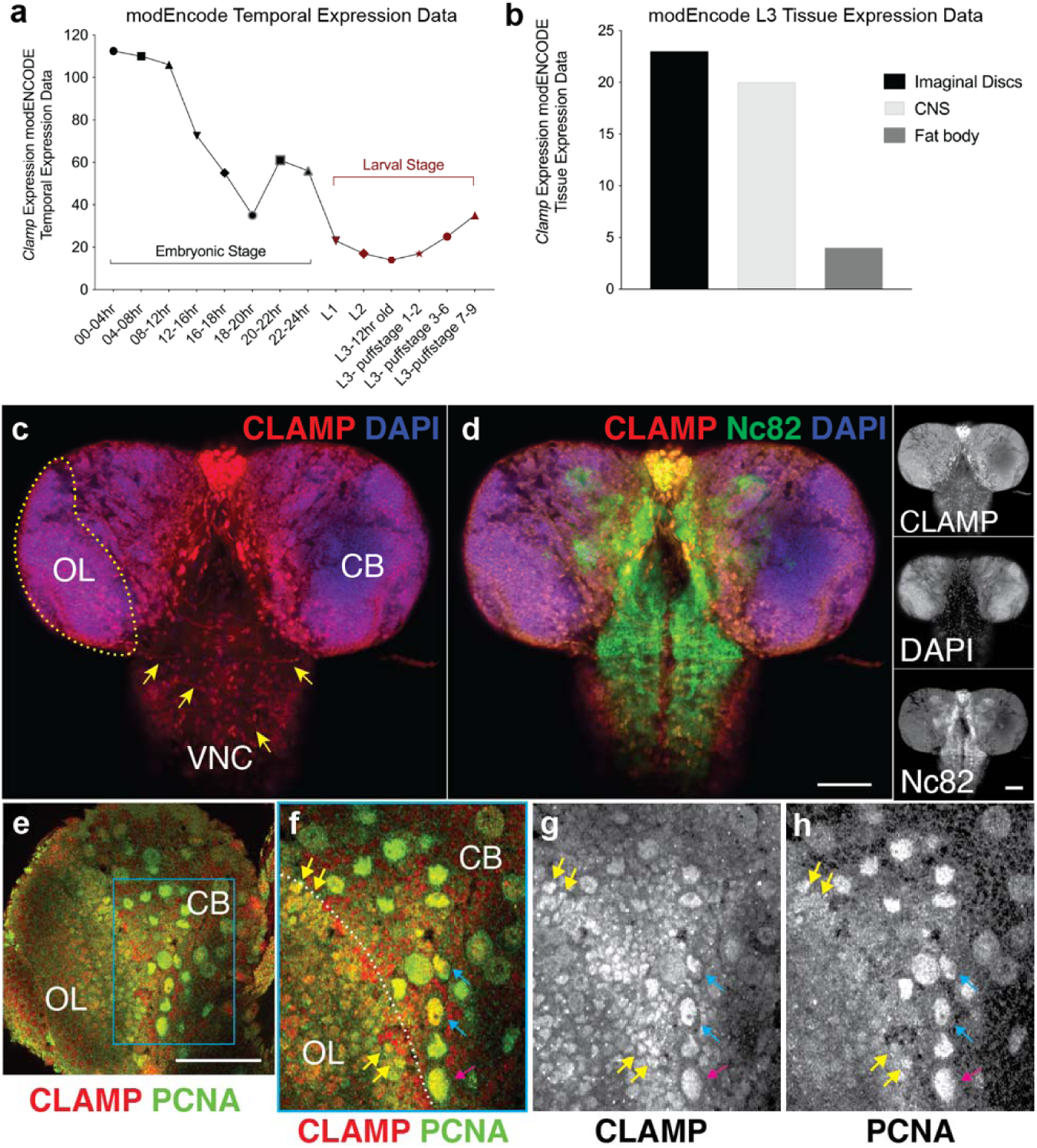
CLAMP is ubiquitously expressed throughout the L3 nervous system and is present in proliferating neuroblasts. (a) Temporal expression data from modEncode (Flybase) reveal that *clamp* expression levels decrease as embryonic development proceeds and as the maternal transcripts are used-up and reach a nadir at 18-20 hours after egg lay, which is then followed by a sharp increase just before embryonic development is completed. During the larval stages *clamp* expression decreases until L3, where it slowly starts to increase again before the pupa stage. (b) Tissue specific L3 expression data, reveal that *clamp* is most highly expressed in the developing imaginal disks, the CNS and the fat body. (c-d) CLAMP is expressed both in the L3 central nervous system, specifically in the optic lobe (OL, yellow dashed outline), central brain, (CB) and in the ventral nerve cord (VNC). CLAMP is present in the nucleus throughout the nervous system as seen by co-staining with DAPI and Nc82, which marks the synaptic processes. In the VNC it is also observed in the neuronal processes (c, arrows). Grayscale panels show individual staining channels. (e-h) Higher resolution images reveal high expression of CLAMP in the developing CB and OL and in proliferating NSCs marked by Proliferating cell nuclear antigen (PCNA). (f-h) Magnification of the boxed area in panel e, reveals that CLAMP is present in CB cycling neuroblasts and mother ganglion cells (magenta and cyan arrows, respectively). Scale bars: 50 μm and 25 μm in j.

To begin to define the role of CLAMP in neural development, we mapped its expression in the developing third instar (L3) central nervous system using immunofluorescence. We found that CLAMP is ubiquitously expressed in the nucleus throughout the larval nervous system, both in cells of the brain and the VNC (Figs. 1c and d). Interestingly, a subset of neuronal processes in the VNC also show protein localization of CLAMP, suggesting that it may be transported to neuronal arborizations (Fig. 1c, axonal processes indicated by arrows). Co-staining of proliferating cell nuclear antigen (PCNA), which labels proliferating cells, revealed that CLAMP is expressed in proliferating NSCs (Figs. 1e-h), both in the central brain and the OL (Figs. 1f-h, cyan and yellow arrows, respectively). CLAMP colocalizes with PCNA both in CB neuroblasts and intermediate precursor ganglion mother cells, as identified by their size difference (Figs. 1f-h, cyan and magenta arrows, respectively). Therefore, we conclude that CLAMP is expressed in proliferating NSCs.

### Loss of CLAMP disrupts larval development

To determine the role of CLAMP in nervous system development, we characterized brain development in *clamp^2^* null mutants that we previously generated using CRISPR (Urban 2017). In these mutants, a 7bp frameshift mutation in the *clamp* gene abolishes expression of CLAMP protein (Fig. 2a, from here on referred to as the *clamp* null). Due to the early embryonic role of CLAMP in the recruitment of the dosage compensation complex in males (Soruco *et al*. 2013b), the majority of male larvae die at the second instar stage. In contrast, female larvae die at the late L3/prepupa stage (Urban 2017). Because of our interest in the essential role of CLAMP, independently of dosage compensation, and due to the small number of male *clamp* null larvae, we focused on female L3 larvae. Comparison of *clamp* null L3 larvae to stage-matched controls (see Materials and Methods) revealed that *clamp* larvae are significantly smaller than controls (Figs. 2a and b). Furthermore, macroscopic examination of the central nervous system (CNS) of *clamp* null larvae revealed that their brains are also significantly smaller than controls (Figs. 2c and d). Western blotting for CLAMP in *clamp* null larvae brains verified the absence of CLAMP protein in *clamp* null L3 brains (Fig. 2e).

**Figure 2.**
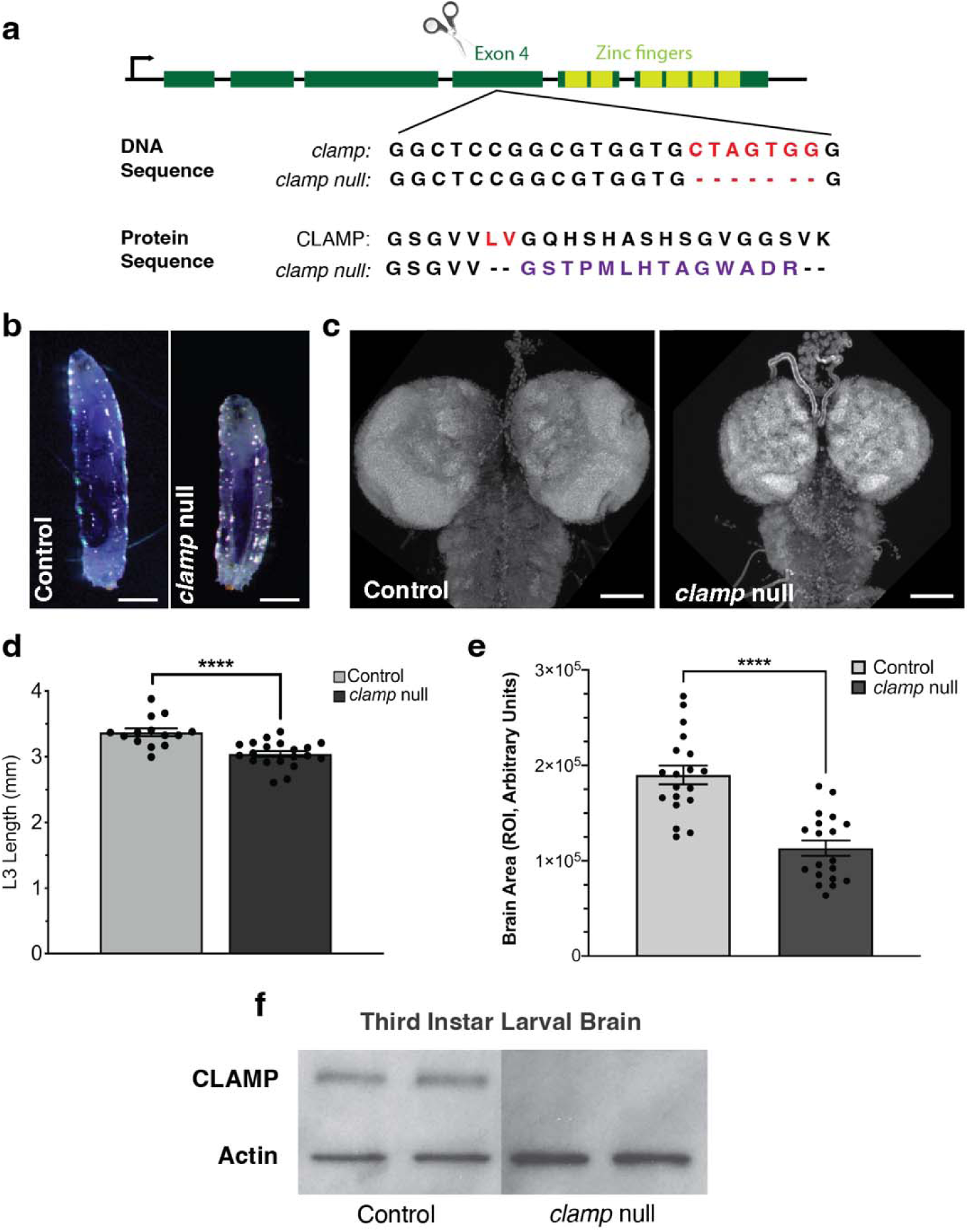
Lack of CLAMP results in growth deficits both in the body and the nervous system (a) Schematic of the CRISPR/Cas9-mutation used to generate the *clamp* null animals. A 7bp deletion was introduced in the fourth exon (dark green boxes) of the *clamp* gene, upstream of the zinc-finger DNA-binding domain, causing a frameshift of the protein sequence (purple) and resulting in an early termination codon. (b, d) Mutant *clamp* null L3 larvae are considerably smaller in size, compared to control L3s (Mean length ± SEM; Control: 3.369 ± 0.06031, n=14; *clamp* null: 3.041 ± 0.04371, n=20; ttest p< 0.0001). (c, e) This reduction is maintained at the CNS level of *clamp* null L3 larvae (Mean normalized brain area ± SEM; Control: 1.001 ± 0.0485, n=20; *clamp* null: 0.601 ± 0.04052, n=20; ttest p< 0.0001). (e) Western blot for CLAMP in samples from L3 brain lysates from control and *clamp* null L3 showing the absence of CLAMP protein (n=2 per condition). Scale bars: 1 mm in a; 50 μm in c.

### Loss of CLAMP disrupts brain development

To evaluate CNS development in *clamp null* mutants, we focused on the mid/late L3 stage (∼120 hours after larval hatching (ALH)), because *clamp* null mutants die during late L3/early pupa stages. To gain temporal insight, we also examined early L3 larvae brains. The overall morphology of the CNS, as observed with DAPI staining, is different between *clamp* nulls and controls (Figs. 3a and e). As neural development progresses through the larval stages, the neuroepithelial (NE) cells of the optic lobe (OL) start to differentiate into neuroblasts (NBs), which will subsequently divide to give rise to neurons and glia (Contreras *et al*. 2019a). We observed that the *clamp* null OL is profoundly reduced in size. Specifically, the *clamp null* OL neuroepithelia at 120 hours ALH resemble the control neuroepithelia at 72 hours ALH (i.e., early L3) (Figs. 3a, b, e and f). Similarly, the *clamp* mutant central brain (CB) of the mid/late L3, has an underdeveloped morphology and resembles the early control CB (Figs. 3f and a).

**Figure 3.**
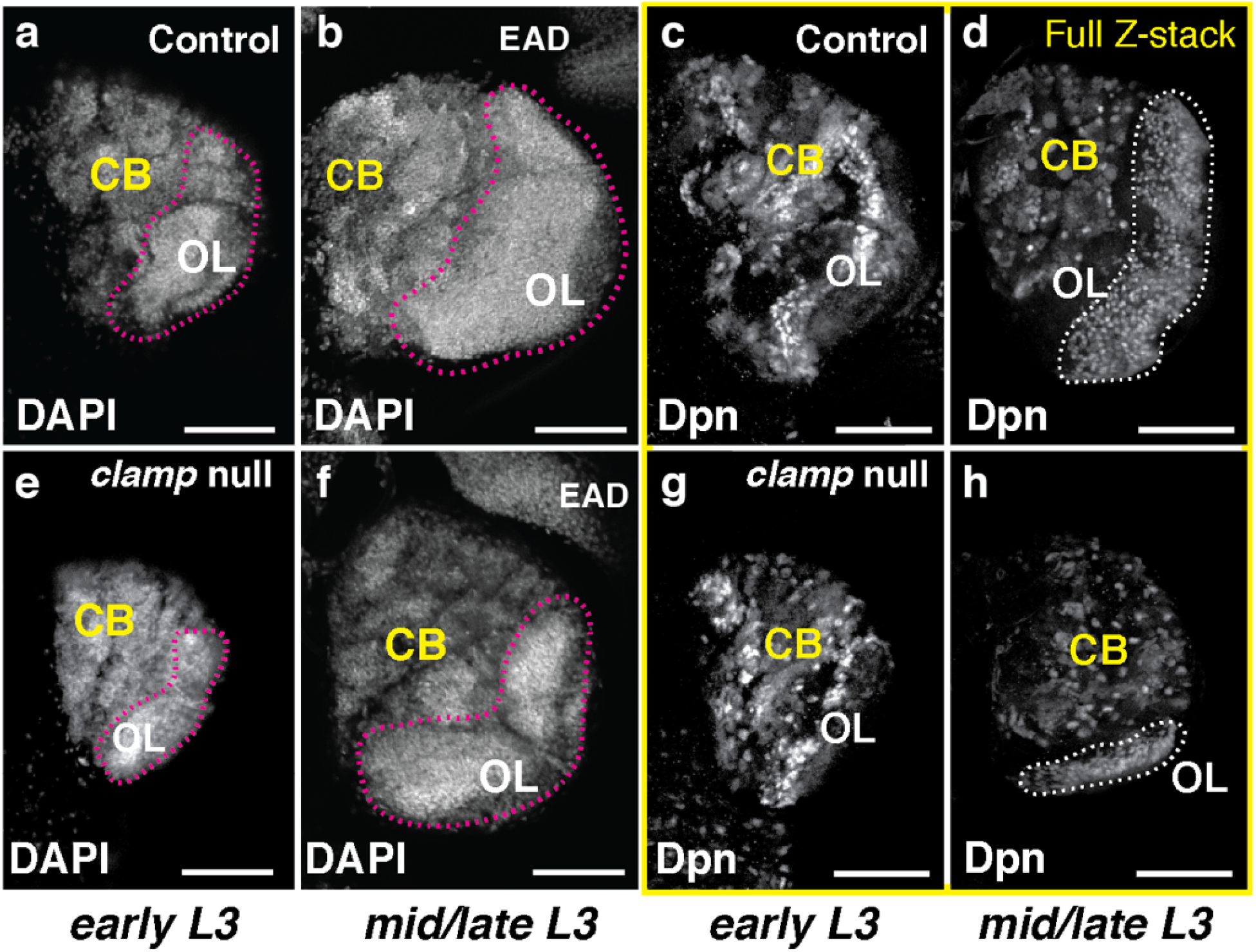
Lack of CLAMP greatly affects brain development and causes defects in optic lobe formation. (a, b and e, f) Brain development in the *clamp* null L3 is severely impaired from the early L3 stage, while even at the mid/late L3 stage, *clamp* null brains resemble early L3 control brains. (c, d and g,h) Full z-stacks of L3 brains showing the distribution of Deadpan (Dpn) -positive NSCs areas, reveals that NSCs development both in the central brain (CB) and the optic lobe (OL) of the null are impaired, starting from the early L3 stage and continuing to the mid/late L3 stage. Especially impaired is the OL development as seen by the reduction in the Dpn+ area indicated with the white dashed outlined areas. EAD: eye antenna disk. Panels a, b, e and f are 3 optical sections whereas c, d, g and h are full z-stacks throughout the brain. Scale bars: 50 μm.

Next, we examined NSCs in *clamp* null and control brains using an antibody against Deadpan (Dpn), which labels all NSCs. Full z-stacks of the Dpn-positive NSCs in early and mid/late L3 brains (Figs. 3c, d, g and h), revealed that in the *clamp* null, the development of NBs, both in the CB and OL, is considerably delayed from the early L3 stage and remains so until the mid/late L3 stage when the animals die. The OL NB population is greatly affected as the area containing Dpn-positive NSCs is much smaller in *clamp* nulls than controls in mid/late L3 brains (Figs 3d and h, dashed outlines). These observations suggest that the deficits observed in L3 *clamp* null brains may be due to an earlier arrest of NBs, resulting in the generation of fewer NSCs. Comparison of early L3 *clamp* heterozygotes (*clamp^+^*^/-^) and *clamp* null brains indicates that a single copy of the *clamp* gene is sufficient to maintain the overall morphology of the developing OL (Supplementary Fig. 1).

### Developmental transcriptomic analysis of clamp null mutants reveals a role for CLAMP in nervous system development

The neurodevelopmental deficits observed in the larval *clamp* null mutant brain suggest that the function of CLAMP initiates earlier than the L3 stage. CLAMP is an ubiquitously-expressed TF and is present throughout the nervous system from early embryonic stages due to the deposition of maternal transcripts into the egg (Rieder *et al*. 2019) (Fig. 1a). Each neurodevelopmental stage is characterized by unique transcriptomic events controlled by specific TF factor networks (Konstantinides *et al*. 2018). Thus, to understand how CLAMP regulates nervous system development on the transcriptomic level, we conducted RNA-sequencing at the following key developmental stages in *clamp* null mutants: A) late stage embryos (16-18 hours), a time point where neurogenesis pauses (Hakes *et al*. 2018) and most NBs enter quiescence (Fig. 4a); and B) mid/late L3 stage brains (Fig. 4b), when neuronal and glial cell production are at their maximum (Avet-Rochex *et al*. 2012). The selection of the specific embryonic time point was based on studies by Haussmann et al. (2008, 2010), where *clamp* was identified as a functional interactor of neural development regulators *embryonic lethal abnormal vision* (*elav*) and *erect wing* (*ewg*) (Robinow and White 1988; Desimone and White 1993; Haussmann *et al*. 2008; Haussmann and Soller 2010a). To control for non-specific transcriptional changes due to other mutations that could be in the genetic background, we used the *CLAMP* rescue line as a control (Urban 2017). In the *CLAMP* rescue line, wild-type *CLAMP* and its regulatory region is inserted into an attP site to restore expression under the control of the endogenous promoter.

**Figure 4.**
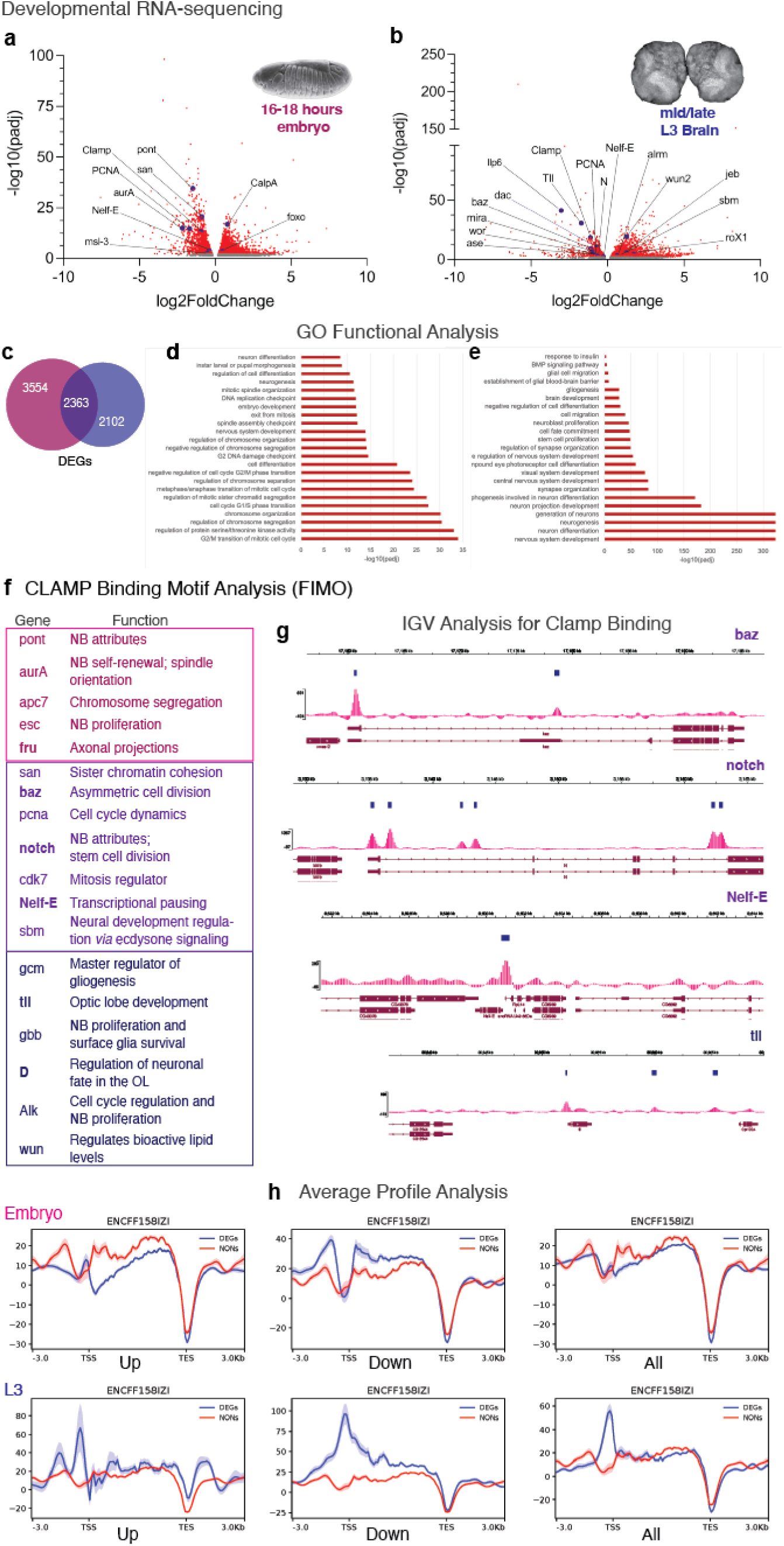
CLAMP is implicated in transcriptomic networks that regulate the proliferation and differentiation of NSCs. (a, b) Experimental workflow for RNA sequencing data generation and DEG meta-analysis. The embryo panel is a scanning electron image of a late stage embryo from (Turner and Mahowald 1977). (c) Venn diagram of DEGs in the *clamp* null embryo and L3. (d,e) GO analysis of the *clamp* null DEGs revealed high enrichment in categories related to cell cycle dynamics and neural development. (f) Find Individual Motif Occurrences (FIMO) analysis of the DEGs for the CLAMP binding motif, revealed that critical regulators of neural development and mitosis are putative targets for CLAMP binding both at the embryonic and L3 stage, (g) while analysis of embryo ChIP-seq data revealed a direct regulation of *Tll, Notch, baz* and *Nelf-E* by CLAMP. (h) Average binding profiles for the upregulated and downregulated DEGs in embryo and L3 *clamp* null. Gene regions were scaled to match positions of the transcription start sites (TSS) and transcription end sites (TES) with 3k base pairs shown on either side of the gene. For comparison in each image, the average read depth is also plotted for 2571 gene regions that show expression in the nervous system but are not differentially expressed in any of these RNA-seq experiments (NONs). The shaded region around each line is the standard error of the mean. The genes downregulated in the absence of clamp show higher average read depth around the TSS, particularly in larvae.

Analysis of our transcriptomic data revealed that many genes were significantly differentially expressed (DEGs) both in the embryo and the L3 brain (padj ≤ 0.05, Figs 4a-c). Interestingly, a large proportion of DEGs were shared between embryo and L3 brain data sets (Fig. 4c). The embryo dataset contains 5917 DEGs (padj ≤ 0.05 and Supplementary Table 1a), while the L3 contains 4465 DEGs (padj ≤ 0.05 and Supplementary Table 2a). Of these DEGs, 2363 are shared between the two stages representing 40% of the embryonic DEGs and 53% of the L3 DEGs. The large number of shared DEGs across time supports the hypothesis that *clamp* regulates a subset of the same genes in both embryonic and L3 stages in addition to stage-specific targets.

GO analysis revealed statistically significant enrichment for developmental/stem cell-related GO categories, including neural development, at both the embryonic and larval developmental stages (Figs. 4d, e). Enrichment was also observed for GO categories involving metabolic pathways. This is interesting as post-embryonic neurogenesis is largely orchestrated by signals regulated by nutrient availability and insulin signaling (Britton and Edgar 1998; Bailey *et al*. 2015; Galagovsky *et al*. 2018).

Functional analysis of the embryonic DEG dataset revealed highly statistically significant enrichment in categories such as cell cycle regulation, cell cycle checkpoints, mitotic cell cycle phase transition, neurogenesis and nervous system development, metabolic processes, synaptic development and signaling (Supplementary Table 1b). We focused our analysis on two major highly enriched GO categories: i). *Cell Cycle Regulation* [GO:0051726, padj: 6.22E-11] and ii). *Nervous System Development* [GO:0007399, padj: 4.32261E-05].

Within the *<u>GO:0051726: Cell Cycle Regulation</u>* category (Supplementary Table 1c), we identified DEGs with specific roles, including factors that regulate the G2/M transition of the mitotic cell cycle, regulation of chromosome segregation, cell cycle G1/S phase transition, negative regulation of nervous system development and of insulin signaling (Fig. 4d).

Meta-analysis of the *<u>GO:0007399: Nervous System development</u>*category (Supplementary Table 1d), revealed strong enrichment in the sub-categories of neurogenesis and central nervous system, visual system development, glial cell development and differentiation (Fig. 4e). Several DEGs critical for neural development and NSC biology were identified, including *Notch* [L2FC: -0.5, padj: 0.01]. Notch is a key regulator of NSC divisions throughout the nervous system (Contreras *et al*. 2018; Hakes and Brand 2019). In the OL, Notch signaling is critical for the equilibrium between proliferation and differentiation of NSCs, especially for the balance between symmetric and asymmetric stem cell divisions, which produce more NSCs or progeny, respectively, in the optic lobe (Egger *et al*. 2010) and neuroblast selection and delamination (Arefin *et al*. 2019). The discovery that loss of CLAMP affects *Notch* expression may provide insight into how CLAMP regulates nervous system development, including the greatly underdeveloped OL of the *clamp* mutants.

Furthermore, we identified additional DEGs implicated in various aspects of NSC function, including: 1) *elav*, *polo, aPKC, par6, pins, aurA* and *baz-* NSC division mode; 2) *PCNA, RFC38, dacapo,* and *Reptin* – NSC fate determination; 3) *CSN1b, CSN4* and *non-stop*, implicated in lamina glia cell migration and optic lobe development (Weake *et al*. 2008). Many of these genes are either regulated by or interact with Notch, suggesting that through the potential regulation of Notch signaling and of the subsequent inter-dependent gene/TF networks, CLAMP is implicated in: 1) cell cycle/asymmetric division; 2) neurogenesis/gliogenesis; and 3) neuronal arborization/synaptogenesis.

### Transcriptomic analysis of shared embryo and L3 *clamp* null DEGs implicates CLAMP in a continuum of embryonic and post-embryonic neurogenesis regulation

The transition from the embryonic to the post-embryonic stage is accompanied by many physiological changes with the most notable being the introduction of feeding which brings about a cascade of metabolic signals inducing activation of quiescent NSCs and neurogenesis. Comparison of the *clamp* null embryo and L3 mRNA-seq results revealed a large number of shared DEGs (2363 DEGs) (Fig. 4b). Meta-analysis of the top two shared GO categories identified genes with key roles in nervous system development, neurogenesis, and mitotic cell cycle, thus suggesting continuing roles for CLAMP in neural development pre- and post-embryonically.

### CLAMP regulates genes controlling NSC attributes, glia development and function

To explore the specific roles of CLAMP in the larval stage, we classified L3 DEGs (Supplementary Table 2b) as downregulated or upregulated in the *clamp* null mutant (downregulated vs. upregulated, Tables 2 and 3, respectively). Downregulated genes are putative targets activated by CLAMP in wild-type animals. These genes could be classified into the following categories: 1) *NSC attributes* (cell-cycle progression, cell polarity, and survival); 2) *control* of *NSC evolution/differentiation*; 3) *regulation of gliogenesis*; 4) *optic lobe development-specific*. In contrast, upregulated genes that are putatively repressed by CLAMP fell into the following categories: 1) *lipid metabolism/promoting mobilization of lipids*; 2) *ecdysone and insulin signaling in glia cells*; 3) *pro-arborization/synapse formation and function*; 4) *pro-neural/suppression of proliferation*. These data suggest that loss of *clamp* leads to deficits in neural stem cell attributes and promotes changes that lead to the precocious acquisition of a neuronal fate at the expense of gliogenesis. Our expression data also suggest that loss of *clamp* may cause concomitant activation of lipid metabolism/mobilization, possibly as a compensatory response from the stem cell-niche. Of great interest, we observed a prominent and strong downregulation of genes orchestrating OL development such as *tll, eya, dac* and *optix* (See Table 2 and Supplementary Tables 2d and Figs. 4b). The downregulation of OL development master regulatory genes in the absence of *clamp* is consistent with the greatly reduced size of the OL in the L3 *clamp* null mutants (see Fig. 3). Collectively, these data are consistent with a critical role for CLAMP in the regulation of OL NSCs.

**Table 1:** Functional interactors of clamp as identified by Haussman et al., 2008.

| Gene Name | log2FC | padj |
| --- | --- | --- |
| *elav* | 0.44 | 0.043 |
| *ewg* | NA | NA |
| *clamp* | -1 | 6.39E-14 |
| *CG12299* | -0.70 | 0.0001 |
| *gro* | -0.88 | 2.45E-06 |
| *CG8924* | -1.12 | 1.15E-10 |
| *JIL-1* | NA | NA |
| *osk* | -4.76 | 1.67E-11 |
| *ac3* | 0.54 | 4.87E-05 |
| *cdc16* | -0.83 | 0.005 |
| *Ibf1* | -0.78 | 0.0002 |
| *Bcl7-like* | -0.93 | 2.92E-11 |

**Table 2:** Down-regulated genes in the clamp null L3 brain.

| *Gene ID* | *Gene Name* | *Log2FC* | *padj* | *Function* |
| --- | --- | --- | --- | --- |
| FBgn0044047 | *Dilp6* | -3.028 | 4.38E-42 | OL development/Glia Neuroblast interactions/Insulin Signaling |
| FBgn0003720 | *tll* | -1.71 | 1.89E-31 | OL development |
| FBgn0031745 | *rau* | -1.373 | 3.42E-18 | Glial cell development/differentiation of neurons and glia in the developing eye. |
| FBgn0014179 | *gcm* | -1.281 | 5.90E-12 | Glial cell development-promotes gliogenesis over neurogenesis |
| FBgn0010433 | *ato* | -1.246 | 3.88E-15 | OL development |
| FBgn0000529 | *bsh* | -1.155 | 1.85E-24 | OL development |
| FBgn0019809 | *gcm2* | -1.154 | 3.97E-20 | Has a redundant role with the product of *gcm* in the differentiation of glial cells and subset of neurons(CHOTARD *et al.* 2005). |
| FBgn0031375 | *erm* | -1.116 | 2.76E-19 | Cell fate regulator during neurogenesis. |
| FBgn0000163 | *baz* | -1.105 | 6.44E-10 | Bazooka forms a complex with par-6 and aPKC. Functions in cell polarization and ACD. |
| FBgn0021776 | *mira* | -0.99 | 3.16E-07 | Neuroblast proliferation. |
| FBgn0000137 | *ase* | -0.972 | 1.54E-05 | Type I NB attributes. |
| FBgn0263512 | *Vsx2* | -0.956 | 9.10E-23 | OL development. |
| FBgn0010382 | *CycE* | -0.951 | 0.00131 | Cell cycle dynamics. |
| FBgn0001983 | *wor* | -0.94 | 3.01E-05 | Worniu, neuroblast attributes. |
| FBgn0000320 | *eya* | -0.909 | 2.05E-08 | OL development /insulin producing cells differentiation. |
| FBgn0000404 | *CycA* | -0.855 | 0.010084 | Cell cycle progression including G2. |
| FBgn0025739 | *pon* | -0.847 | 3.66E-07 | Contributes to asymmetric localization of numb and the subsequent suppression of Notch signaling. |
| FBgn0002973 | *numb* | -0.845 | 1.67E-15 | Acts via Notch signaling, asymmetric division control. |
| FBgn0005655 | *PCNA* | -0.824 | 0.00019 | Cell cycle dynamics. |
| FBgn0011674 | *insc* | -0.788 | 1.44E-06 | Spindle orientation; asymmetric divisions. |
| FBgn0037913 | *fabp* | -0.72 | 1.85E-06 | Expressed in lipid droplet-forming glial cells. |
| FBgn0013469 | *klu* | -0.713 | 0.00013 | Klumpfuss (*klu*) is a temporal fate determination. |
| FBgn0005677 | *dac* | -0.707 | 4.97E-11 | OL development. |
| FBgn0010109 | *dpn* | -0.665 | 0.00245 | Neuroblast self-renewal -interacts with *klu* and *erm*. |
| FBgn0004647 | *Notch* | -0.642 | 0.0018 | OL development/Stem cell proliferation. |
| FBgn0000411 | *Dichaete (D)* | -0.634 | 2.56E-08 | Nervous system development and OL development. |
| FBgn0044324 | *Chro* | -0.61 | 0.00021 | Neuroblast proliferation. Functions downstream of InR/PI3K/TOR signaling and regulates expression of quiescence-inducing Pros to permit NSC reactivation. |
| FBgn0005427 | *ewg* | -0.533 | 1.06E-06 | Neurogenesis/Synaptogenesis. |
| FBgn0040080 | *pins* | -0.52 | 0.008985 | Partner of Inscutable; involved in ACD and establishment of cell polarity. |

**Table 3:** UP-regulated genes in the clamp null L3 brain.

| *Gene ID* | *Gene Name* | *Log2Fc* | *padj* | *Function* |
| --- | --- | --- | --- | --- |
| FBgn0000014 | abd-A | 2.983 | 0.00019 | Controls the end of neuroblast proliferation. |
| FBgn0025879 | Timp | 2.299 | 9.57E-23 | Coordinates synaptogenesis |
| FBgn0001258 | Ldh | 2.353 | 2.43E-22 | Lipid metabolism. |
| FBgn0003117 | pnr | 2.106 | 0.008812 | Activator of *proneural* achaete-scute complex genes. |
| FBgn0011746 | ana | 1.831 | 3.97E-20 | Encodes a secreted glycoprotein that is expressed in glia and at lower levels in neuroblasts, <u>suppresses neuroblast proliferation</u>. |
| FBgn0032006 | Pvr | 1.742 | 8.56E-22 | Glia development /neuroblast proliferation. |
| FBgn0016032 | lbm | 1.576 | 3.84E-21 | facilitates synapse assembly |
| FBgn0037989 | ATP8B | 1.664 | 2.18E-20 | Flippase, translocates phospholipids from one side of a membrane to the other forming new membrane structures, especially vesicles. |
| FBgn0031461 | daw | 1.502 | 2.82E-05 | Regulates insulin release from IPC neurons. |
| FBgn0041087 | wun2 | 1.249 | 3.17E-20 | Encodes a lipid phosphate phosphatase; expressed in astrocytes. |
| FBgn0086677 | jeb | 1.224 | 1.37E-06 | Functions as signaling ligand for the product of <u>Alk</u>. |
| FBgn0066101 | LpR1 | 0.899 | 3.91E-21 | Cellular uptake of neutral lipids from the circulating protein encoded by apolpp. |
| FBgn0000416 | Sap-r | 0.889 | 2.23E-29 | Lysosomal Lipid homeostasis. |
| FBgn0000546 | EcR | 0.847 | 6.72E-06 | Ecdysone Receptor. |
| FBgn0040505 | Alk | 0.692 | 0.00014 | Employs Ras/ERK and PI3K signalling pathways, functions in multiple contexts including synaptic signaling. |
| FBgn0001297 | kay | 0.699 | 7.62E-12 | Specifically expressed in glia. |
| FBgn0039332 | alrm | 0.635 | 0.00038 | Glial related. |
| FBgn0000370 | crc | 0.61 | 3.83E-17 | Coactivator of the product EcR, promotes expression the molting peptide hormone encoded by ETH. |
| FBgn0016078 | wun | 0.558 | 2.85E-07 | Lipid phosphate phosphatase. |
| FBgn0013342 | nSyb | 0.506 | 3.24E-05 | Synaptic signaling. |
| FBgn0030574 | sbm | 0.504 | 0.002373 | L-type amino acid transporter expressed in glia; essential for proper timing of development and brain growth (GALAGOVSKY *et al.* 2018). |

### CLAMP binds to chromatin at major neurodevelopment genes

To evaluate the possibility of direct regulation of DEGs by CLAMP, we mapped the presence of CLAMP binding sites using FIMO analysis (Grant *et al*. 2011). Previously characterized CLAMP binding motifs were used for this analysis (Kaye *et al*. 2018). Interestingly, genes that contain putative CLAMP binding sites and are dysregulated in *clamp* nulls both in the embryonic and L3 stages include the critical proliferation/mitosis regulator *Notch*; *bazooka (baz); separation anxiety* (*san)*, a N-terminal acetyltransferase required for mitotic chromosome segregation (Ribeiro *et al*. 2016); *cyclin dependent kinase 7* (*cdk7*), vital for mitosis (Larochelle *et al*. 1998); and *PCNA*. CLAMP also interacts with chromatin at *sobremesa* (*sbm*), an SLC7 amino acid transporter expressed in glia that regulates brain development *via* ecdysone signaling (Galagovsky *et al*. 2018), and *Negative elongation factor E* (*nelf-E*), which is a known interactor of CLAMP at the protein level (Urban 2017) (Fig. 4h, Embryo and L3 panels).

CLAMP binding sites are present across stage-specific neurodevelopmental genes. Embryonic neuronal genes with CLAMP binding sites include *anaphase promoting complex subunit 7* (*APC7*)*; pontin (pont),* a component of the TIP60 complex which regulates NSC attributes*; aurora A (aurA)* a kinase regulating asymmetric protein localization and thus NSC proliferation (Lee *et al*. 2006)*; extra sex combs (esc)*, a component of the PRC2 complex which regulates NSC proliferation and neural development (Curt *et al*. 2019); and *fruitless (fru)*, implicated in the formation of axonal projections in the embryonic CNS (Song *et al*. 2002) (Fig. 4e, Embryo panel). Notably given its role in OL development, neuronal genes expressed during the L3 stage with CLAMP binding motifs include tailless (*tll),* a key regulator of optic lobe development and NSC function; *gcm,* an essential regulator of glial development*; glass bottom boat* (*gbb*) which regulates both NB proliferation and surface glia survival; *Dichaete* (*D*), a TF regulating neuronal fate in the OL; and *anaplastic lymphoma kinase (Alk)*, which is implicated in cell cycle regulation and NSC proliferation (Fig. 4f, L3 panel).

To examine and validate which of the FIMO-identified CLAMP binding targets have been experimentally confirmed for CLAMP binding, and evaluate the basis for direct CLAMP regulation, we mined available embryonic 0-24 hour CLAMP ChIP-seq data generated by modENCODE (Consortium *et al*. 2010) and L3 whole larval data (Kuzu *et al*. 2016) using the Integrative Genomics Viewer (IGV) (Robinson *et al*. 2011). This analysis revealed that CLAMP directly binds to *baz and Nelf-E,* the latter of which serves as a positive control to our results as an established direct target of CLAMP (Urban 2017) (Fig. 4g). Importantly, CLAMP directly binds to and positively regulates both *Notch* and *Tll* expression (Fig. 4g and see Fig. 4b), suggesting that the *clamp* null mutant OL development defects may be due to downregulation of *Notch* and *tll* expression in the absence of CLAMP.

To further assess direct regulation by CLAMP, we quantified CLAMP binding to DEGs in both our data sets compared to a control gene set of random neuronally expressed genes. In support of a direct regulatory role for CLAMP at a subset of its target genes, we found that genes that are downregulated in *clamp* nulls show higher average read depth around the TSS (Fig. 4h) and have a greater binding frequency than at control genes, particularly in larvae (Supplementary Table Y).

### CLAMP promotes NSC proliferation and prevents precocious neurogenesis

Our transcriptomic, motif and ChIP-seq analyses demonstrate that CLAMP is exerting direct control on neural development and fate determination by directly regulating the transcription of key effectors *Notch* and *tll*, (Figs.4f, g). Concurrently, in the absence of CLAMP we observed a marked decrease in genes related to glial-development and cell cycle dynamics (Table 2), processes in which *Notch* and *tll* are integral, particularly in the optic lobe where they regulate neuronal versus glial identity and stemness potential (Guillermin *et al*. 2015; Ren *et al*. 2018). Therefore, we hypothesized that lack of CLAMP disturbs the balance between neuronal and glia formation through the L3 stage, by impinging neural stem cell proliferation, especially in the OL.

To test our hypothesis, we evaluated mid-late L3 brains for proliferating NSCs using the S-phase marker EdU, the NSC marker Dpn and a quantification approach introduced by Kanai et al.(Kanai *et al*. 2018) in which the superficial layers of the brain are evaluated. We chose this approach due to the importance of superficial glia in the overall biology of neuroblasts and used it throughout our study. Consistent with our initial observations (Fig. 3), the OL in *clamp* null larvae brains was dramatically reduced in size compared to controls, while both the CB and OL contained fewer proliferating NSCs (EdU^+^Dpn^+^, Figs. 5a and c and Supplementary Figs. 2a, b). Quantification of total proliferating NSCs (EdU+) from the superficial layers of the brain (Kanai *et al*. 2018), showed a dramatic ∼90% reduction compared to controls (EdU^+^Dpn^+^ Fig. 5e). Moreover, we observe an overall reduced number of proliferating cells compared to controls, suggesting a reduced proliferation of stem cells and/or change in cell cycle dynamics in the absence of CLAMP (EdU+ Fig. 5e).

**Figure 5.**
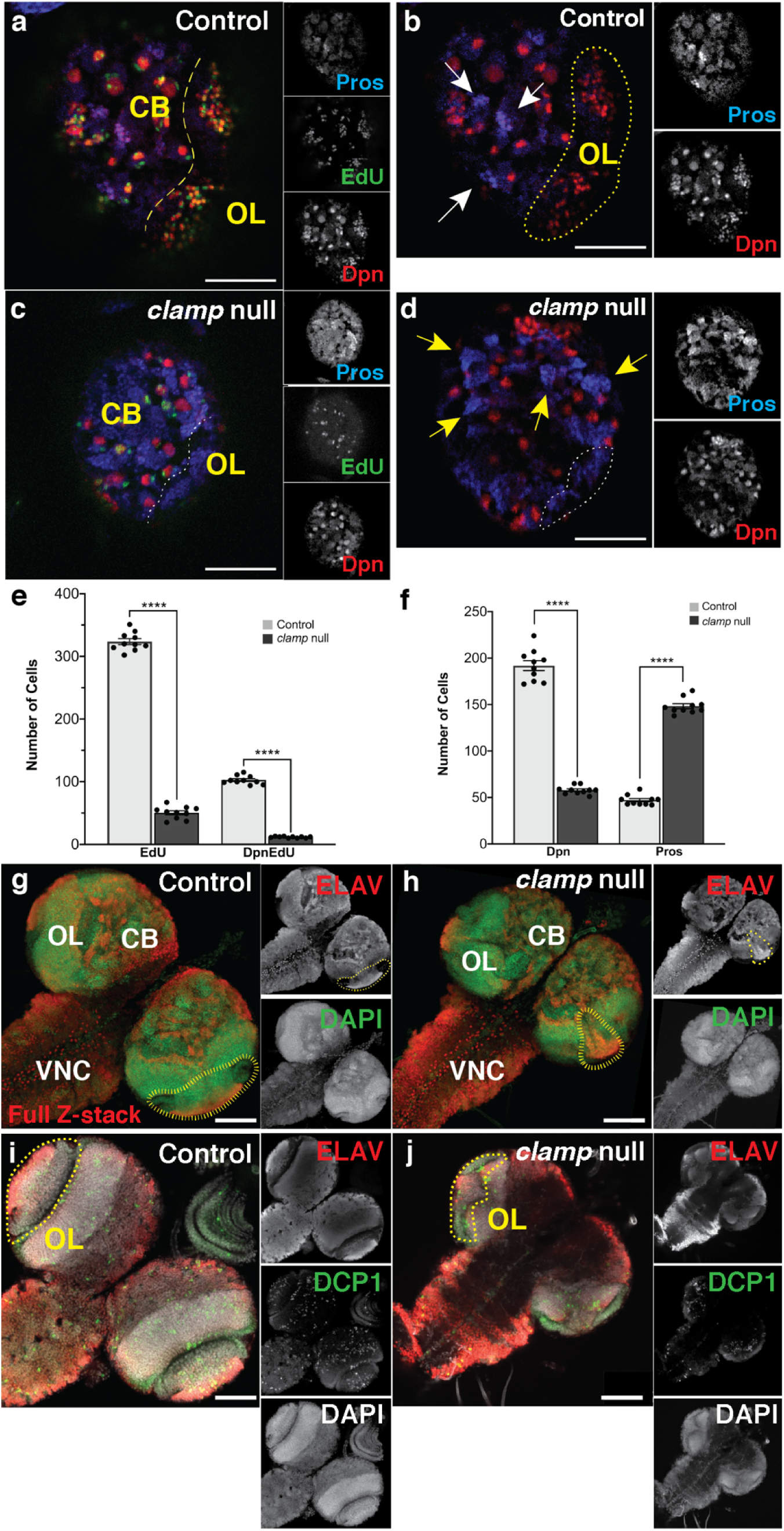
Lack of CLAMP impairs neural stem cell proliferation and promotes precocious differentiation. (a-d) Proliferation of NSCs as seen by EdU labeling, is highly reduced in the *clamp* null brain, with the most evident impact being on OL NSCs (panels a and c, dashed lines indicate the OL). In parallel, nuclear Pros+ clusters indicate acquisition of neuronal fate, are more evident in the *clamp* null brains (panels b and d, arrows). (e) Both the total number of proliferating stem cells (EdU^+^ Mean ± SEM, Control: 323.6 ± 4.679; *clamp* null: 50.40 ± 3.174; p<0.0001; n=10) and of proliferating Dpn^+^ NSCs is reduced (Dpn^+^EdU^+^ Mean ± SEM; Control 102.6 ± 2.141; *clamp* null 11.50 ± 0.3727; n=10). (f) The reduction in total Dpn+ cells (Dpn^+^ Mean ± SEM, Control: 191.9 ± 5.313; *clamp* null: 57.90 ± 1.402; p <0.0001; n = 10), is accompanied by a respective increase in Pros+ cells in the *clamp* null brains (Pros^+^ Mean ± SEM, Control: 47.1 ± 1.716; *clamp* null: 148.3 ± 2.629; p <0.0001; n = 10). (g, h) Additionally, despite having an increase in the neuronal marker Elav in the *clamp* null brains, the OL development is impaired compared to controls (outlines). (i, j) Apoptotic cells marked with DCP1 are more in the OL of the null compared to a more ubiquitous distribution in the control. CB, central brain; OL, optic lobe; VNC: ventral nerve cord. Scale bars: a-d, 100 μm; g and h, 50 μm.

The TF Prospero (Pros) acts as a master regulator driving the decision of stem cells to either self-renew or differentiate (Choksi *et al*. 2006), with nuclear Pros signaling the initiation of neuronal differentiation (Knoblich and Jan 1995; Choksi *et al*. 2006). Thus, to evaluate changes in neuronal production in the *clamp* null brain we performed Pros immunostaining. We found that the *clamp* null brain contained more clusters of nuclear Pros immunostaining than matched controls, thus indicating a rapid change in NSC cell fate attributes (Colonques *et al*. 2011) (Figs. 5b and d, arrows indicate Pros+ clusters). Interestingly, while the total number of NSCs in the *clamp* null brain was reduced by approximately 70% (Dpn^+^, Fig. 5f), Pros+ cell numbers were increased by approximately 70% (Pros^+^, Fig. 5f), indicating that stem cells are acquiring a precocious intermediate precursor (ganglion mother cell)/neuronal fate. The reduction in NSCs is particularly evident in the OL, where very few Dpn+ cells were observed, contrary to a widespread presence of Pros+ cells (Figs. 5a-d, outlines in Figs.5b and d indicate the OL). Pros, acts a transcriptional repressor of self-renewal genes and induces differentiation (Choksi *et al*. 2006; Lai and Doe 2014). While a significantly higher number of cells have nuclear Pros (Pros+, Sup.Fig. 2c) in the *clamp* null brains, most of these cells have not incorporated EdU (Pros+EdU-, Sup.Fig. 2c). In combination with the finding that in the *clamp* null brain, a lower number of cells incorporate EdU within the pulse window, and thus a smaller number of NSCs are actively proliferating (EdU^+^Dpn^+^ Fig. 5e), the Pros immunostaining suggests that in the absence of CLAMP, NSCs undergo fewer cell division cycles and/or precociously exit the cell cycle to acquire a differentiated fate.

To further evaluate changes in the neuronal pool of cells, we performed Elav staining to label neurons (Robinow and White 1988), which revealed that *clamp* null brains contain a higher density of Elav^+^ clusters compared to controls (Figs. 5g and h). However, even though the overall Elav+ cluster density increased, the Elav^+^ zone in the OL was reduced compared to controls (Figs. 5g and h, yellow outlines). Immunostaining with the apoptotic marker Dcp1 revealed increased and specific presence of Dcp1^+^ nuclei in the OL in *clamp* nulls, compared to the controls which instead had more widespread apoptosis (Figs. 5i and j). Moreover, in more dorsal optical sections large Dcp1+ clusters were visible in the medulla and lamina area (Supplementary Figs. 3a and b). These results suggest that in the absence of CLAMP a pro-differentiation pathway is activated in stem cells at the expense of their self-renewal which especially impacts the OL. In addition, lack of CLAMP results in increased apoptosis, especially in the OL.

### Expression of the optic lobe development master regulator Tailless, is severely reduced in the clamp null

Our results thus far revealed a direct control of the key OL development gene *tailless* by CLAMP, manifesting as downregulation of *tll expression* (Fig. 6). Interestingly, in *tll* knockdown animals, the OL is severely underdeveloped or absent, OL neurogenesis is affected and OL neuroblasts undergo increased apoptosis, similarly to the *clamp* null OLs (Guillermin *et al*. 2015), reminiscent to the severely underdeveloped OL phenotype of the *clamp* null. To examine whether the *clamp* null OL phenotype can be attributed to a loss of the Tll protein, we performed immunofluorescence for Tll. This approach revealed a consistency between the dramatic reduction in the *tll* transcript identified by mRNA-seq and the protein, with the latter being almost absent in the OL neuroepithelium of *clamp* nulls (Fig. 6d). Additionally, *tll* expression is required for the survival of OL NSCs, since its absence leads to apoptosis (Guillermin *et al*. 2015), which is in agreement with the increased and specific cell-death we observed in the OL region (Figs. 5i and j and Supplementary Figs. 3a and b).

**Figure 6.**
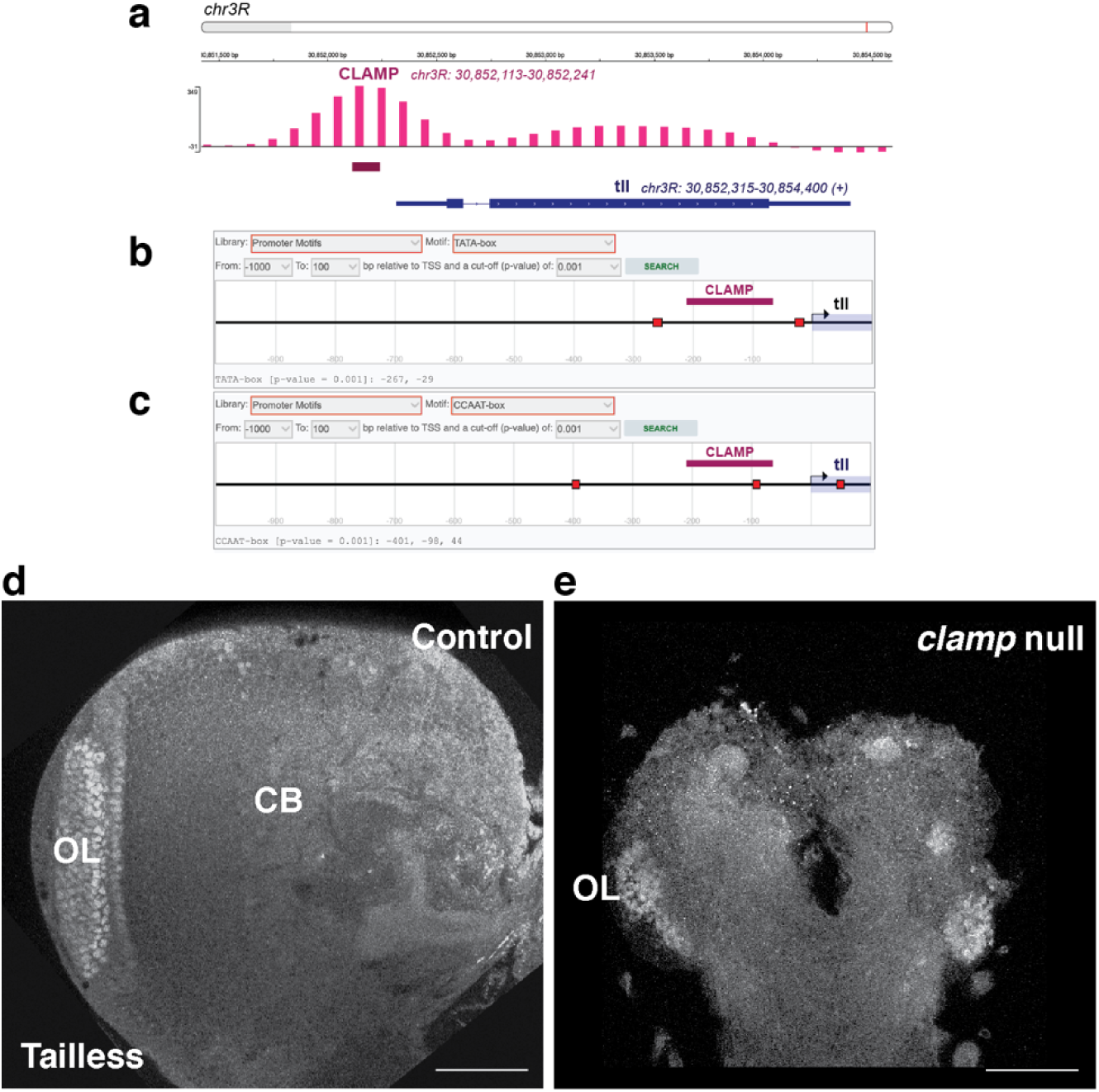
Direct regulation of *Tailless* by CLAMP causes an almost complete absence of its expression in the OL. (a-c) Analysis of embryo ChIP-seq data reveal that CLAMP directly binds to the *Tailless* gene in putative promoter areas. (d,e). Expression of the Tailless protein is almost absent in the clamp null optic (OL) lobe which is severely underdeveloped. Central brain (CB). Scale bars, 50 μm.

### Lack of CLAMP results in reduction in number of surface glia

OL development is highly dependent on glia-neuron interactions because glia form a supportive niche for neural stem cells to function (Pereanu *et al*. 2005; Ramon-cañellas *et al*. 2018). Furthermore, neuroepithelial cells of the OL give rise to hundreds of precursor/progenitor cells of different lineages, including neuroblasts and glial progenitor cells. For the lineage diversification to take place an intricate temporal expression of fate-determining TFs occurs in the precursor cells, including *Notch* and *Tll,* both of which are under direct CLAMP control (Fig. 4g). In OL medulla NBs, Tll is expressed in the oldest NBs which go on to generate neuronal progeny or express *glia cells missing (gcm)* and gradually turn off *Tll* to express *Repo* as glial cells (Li *et al*. 2013).

Our transcriptomic analysis supports the hypothesis that lack of CLAMP promotes neuronal differentiation at the expense of gliogenesis from the late embryo stage. Therefore, the defects observed in OL development of the L3 *clamp* null could be attributed to a combination of deficits in gliogenesis and precocious neurogenesis as a result of altered *Tll and/or Notch* expression, since *Notch* is known to regulate *Tll* dynamics to coordinate neural specification in the larval brain and the optic lobe (Mishra *et al*. 2018). Surface glia (SG) are crucial for NB proliferation (Kanai *et al*. 2018), an interplay which involves niche and autocrine signaling involving the BMP-ligand Glass bottom boat (Gbb). Interestingly, the *gbb* gene contains CLAMP-binding motifs (Fig. 4f) and is downregulated in the *clamp* null L3 brains (Supplementary Table 2d). Furthermore, we observed enrichment of the BMP-signaling GO category in the *clamp* null (Fig. 4d). Therefore, a loss of the dynamic interplay between SG and NSCs resulting from the altered expression of key fate-determining TFs *Tll* and *Notch* in the absence of CLAMP could also contribute to the deficits in the *clamp* null brain.

To test this hypothesis, we first examined whether a change in SG numbers is seen in the *clamp* null. We used the glia-specific marker Repo (Xiong *et al*. 1994b; Yuasa *et al*. 2003) to mark glial cells, and quantified SG in L3 brains using the approach by Avet-Rochex et al. (2012) (Avet-rochex *et al*. 2012). This revealed a ∼25 % reduction in SG number in the *clamp* null, compared to the control (Figs. 7a-d). Additionally, the distribution and numbers of glial cells throughout the brain (Supplementary Figs. 3c and d) and especially in the OL and the lobula plug was altered in terms of topological organization (Supplementary Figs. 3e and f). Overall, these data suggest that CLAMP regulates gliogenesis and glia distribution especially in the OL.

**Figure 7.**
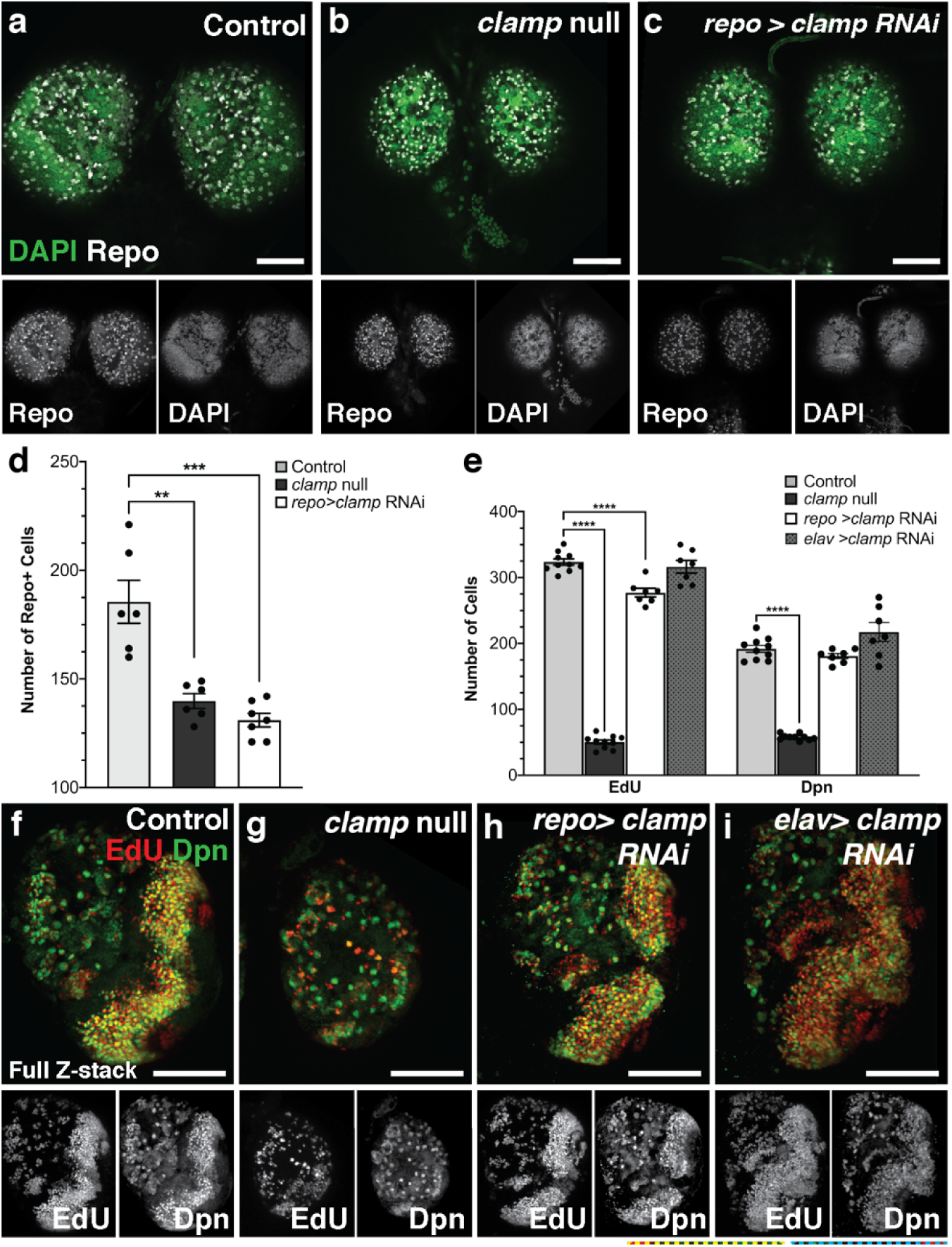
Glial-CLAMP surface glia generation and NSC proliferation. (a-e) Surface glia (SG) is reduced by ∼25% in the *clamp* null brain compared to controls (% SG Repo^+^ ± SEM, *Control*: 99.98 ± 5.328; *clamp* null: 75.37 ± 1.843; n=6) whereas, (e-i) proliferating NSCs (EdU^+^ ± SEM, *Control:* 323.6 ± *4.679; clamp* null: 50.40 ± *3.174*) and NSCs (Dpn^+^ ± SEM, *Control:* 191.9 ± *5.313; clamp* null: 57.90 ± *1.402*) are also reduced in numbers. Glial-specific *clamp* knock-down (*repo>clamp RNAi*) promotes a similar reduction in SG numbers ((% SG Repo^+^ ± SEM, *repo>clamp RNAi* 131.0 ± 3.162) and promotes a reduction in proliferating NSCs (*repo>clamp RNAi* EdU^+^ ± SEM: 277 ± 6.572; Dpn^+^ ± SEM: 181 ± 3.964), but to a lesser extend as the one seen in the *clamp* null. Neuronal-specific clamp KD (*elav>clamp RNAi*) does not affect NSC proliferation and numbers NSCs (*elav>clamp RNAi* EdU^+^ ± SEM: 316. ± 9.519; Dpn^+^ ± SEM: 217 ± 14.45). Scale bars 100 μm.

To determine if the effect on SG number was due to the lack of CLAMP in glia specifically, we used the GAL4-UAS system to induce *clamp*-RNAi knock down (KD) using the glia-specific driver, *repo-GAL4 (Xiong et al. 1994a)*. Interestingly, we find that in the *repo>clamp RNAi* brains, the number of SG was reduced to a similar extent as in the *clamp* nulls (Repo+ cells, ∼30% reduction, Figs. 7a-d). To determine whether glial-specific or neuronal-specific *clamp* depletion has differential effects on the proliferation (EdU+) and the number of NSCs (Dpn+ cells), we also performed neuronal *clamp* KD by crossing to an *elav-GAL4* fly line (Lin and goodman 1994). Comparison with the data we already obtained from the *clamp* null and control brains (see Figs.5e and f) revealed that *clamp* KD in neuronal lineages did not result in significant changes either in EdU+ or Dpn+ cells, while glial-specific *clamp* KD caused a reduction in the number of total proliferating cells (EdU+ cells, ∼15% reduction compared to control). Thus, while glial *clamp* KD results in the same amount of reduction in Repo+ SG number as in the *clamp* null, the effect on proliferating NSCs is smaller compared to the null. This suggests that downregulation of glial CLAMP mainly affects the proliferation of glia-producing NSCs likely *via* both cell autonomous mechanisms (e.g., reduction in Dpn+ gliogenic NSCs) and *via* non-cell autonomous mechanisms (regulation of proliferation by niche glia), although it does not recapitulate the full phenotype seen in the *clamp* null leaving room for additional mechanisms.

In summary, our results suggest that CLAMP controls NSC stemness, proliferation and lineage output, especially in the OL, by directly regulating the expression of key TFs such as *Notch* and *Tailless,* and by affecting neural stem cell niche homeostasis. Ultimately, lack of CLAMP leads to imbalance in neuronal over glial generation and leads to precocious differentiation of neural stem cells.

## Discussion

The understanding of the gene regulatory mechanisms by which neural stem cells function to ultimately build the brain is critical to provide insight into normal development and disease states. Signaling relays originating from the stem niche play an instructive role in the neurogenetic process and highlight the fact that the environmental context acts synergistically with intrinsic developmental signals (Scadden 2006; Speder and brand 2018). In this study, we have defined a new role for the essential transcription factor CLAMP, in nervous system development, NSC-niche interactions, with more profound effects on optic lobe development. We show that CLAMP binds directly to *Notch* and *Tailless* and positively regulates their expression. Through the control of these master neurogenesis regulators, CLAMP impacts NSC stemness, proliferation and progeny output. The multilayered control of CLAMP over NSC function is evidenced in *clamp* null brains, where the OL is severely underdeveloped, and niche glial cell generation is reduced in favor of precocious neurogenesis. Furthermore, lack of *clamp* affects the expression of key NSC proliferation/division genes such as *baz*. In summary, our study reveals that CLAMP regulates brain development *via* cell-intrinsic and NSC niche-dependent mechanisms that involve niche glia.

The formation of the OL is profoundly impaired in *clamp* null mutants, a phenotype accompanied by reduction of EdU^+^ NSCs (Figs. 5a-d). We also observed increased numbers of Pros^+^ cells in the null OL (Figs. 5b and d, outlined area) and throughout the brain, consistent with CLAMP promoting a neurogenic over a gliogenic fate. While glial *clamp* KD results in a partial reduction in the number of proliferating NSCs (*repo*>*Clamp RNAi*: EdU^+^, Fig. 6e), it does not affect the number of Dpn^+^ NSCs i.e., neuroblasts. These results suggest an effect on the Dpn^-^ progeny of the NSCs, such as the ganglion mother cells (GMCs) and common progenitor cells (CPCs) of the OL. This is an intriguing scenario as CPCs divide asymmetrically to generate Lawf precursor cells and glial precursor cells (GPCs) since glial *clamp* KD resulted in approximately the same reduction in Repo+ glial cells, as seen in the *clamp* null. Although, Pros+ cells were not counted in the glial *clamp* KD brains, evaluation of the mean signal intensity of the Pros channel suggested an increased signal compared to the control (not shown), suggesting that lack of CLAMP alters the balance between neuronal and glial progenies at the precursor level.

CLAMPs’ specific role in OL development is evidenced by the fact that CLAMP directly binds and positively regulates *Notch* and *tll* expression. Since we could only obtain functional Tailless antibodies (Notch antibodies tested for IF did not work- results not shown), we were able to confirm downregulation of the Tailless protein expression in the *clamp* null OL (Fig. 4). Furthermore, other transcription factors critical for OL neurogenesis such as *eya, dac* and *optix* are downregulated in the *clamp* null mutant brains (see Table 2).

The OL contains a special population of NSCs, the common progenitor cells (CPCs), which generate both glia precursor cells (GPCs) and neurons (Lawfs), under the control of Notch signaling (Chen *et al*. 2016). Moreover, Notch is a key regulator which maintains the balance between symmetric/self-renewing and asymmetric NSCs divisions; loss of Notch function promotes precocious differentiation and premature neurogenesis (Egger *et al*. 2010). In the absence of CLAMP’s direct regulation (Figs. 4f and g)*, Notch* is at least two-fold downregulated both in the embryonic and L3 stages (Supplementary Table 1a and Table 2). Overall, our data support a model in which CLAMP is upstream of *Notch* and *Tll* signaling to regulate NSC proliferation and differentiation in the OL.

Neural development and NSC function are intricately linked to niche glia and depend on support and nutrition from surface glia *via* Gbb-Dlp signaling, which sustains NSC proliferation and survival but also ensures the survival of surface glia (Kanai *et al*. 2018). Thus, there is an interdependency between NSCs and niche glia, ensuring overall homeostasis in the stem cell niche. In *clamp* null and *repo*>*clamp* RNAi larvae brains, surface glia numbers are reduced to the same extent indicating that lack of glial CLAMP affects gliogenesis (Fig. 7d). Furthermore, the glial master gene, *gcm,* contains CLAMP binding motifs and is significantly downregulated in the *clamp* null brain, indicating that CLAMP may directly regulate gliogenesis through *gcm*-related mechanisms (Fig. 4f). Supportive of a glial role for CLAMP is the fact that glial-specific *clamp* KD by *repo*-GAL4, resulted in specific reduction of SG cells, to the same extent seen in the *clamp* null. In contrast, a neuronal specific *clamp* KD (*elav*> *clamp* RNAi) did not affect NSC proliferation, number or glial cell number (Fig. 6). A small reduction in cycling cells (∼15%, EdU+) was observed in *repo*>*clamp* RNAi conditions compared to a massive 85% reduction in the *clamp* null brain. This difference can be attributed to the fact that in the *repo>clamp* RNAi conditions, the reduction in *clamp* levels will only take place after *repo* expression is established during terminal differentiation of glial cells, and through direct induction by Gcm (Yuasa *et al*. 2003).

In the future, it will be interesting to determine whether this reduction in glial cell number can be attributed specifically to glia-producing cells in which *gcm* activity is still high and indicative of their proliferation status. In the *clamp* null mutants, *clamp* is depleted before *repo* starts to be expressed and thus may result in a greater impact on NSC proliferation and neuronal production in the OL because both *gcm* and *gcm2* regulate glial and neuronal development in the OL (Chotard *et al*. 2005). Thus, it is possible if we used *gcm*-GAL4 to drive *clamp* RNAi at earlier stages, we would unmask a greater effect both on NSC proliferation and NSC number.

Our results indicate that CLAMP is important both for maintaining the proliferation of NSCs by acting in a cell-type specific manner (i.e, glia vs. NSCs) and in a cell-intrinsic mechanism that implicates direct regulation fate determinants *Tailless* and *Notch,* especially in the OL. Our results are further strengthened by the recent study of Coyne et al. (Coyne *et al*. 2025), documenting the specific role of CLAMP neurogenesis, since CLAMP binding motifs were shown to be prevalent in neuronal enhancers and regulating the balance of NSC proliferation and premature activation. Overall, here we have shown that CLAMP is an important regulator of NSC properties and of the stem cell niche.

## Acknowledgements

We thank Dr. Edward A. Kravitz for providing MAT laboratory space and contributing to supply costs for completion of this study, Dr. Yick-Bun Chan for experimental support and assistance with DEG analysis and Dr. Yiannis Savva for experimental support. We would also like to thank Dr. Séverine Trannoy for critical reading of this manuscript. We thank the Harvard Medical School Neurobiology Department and the Neurobiology Imaging Facility for consultation and instrument availability that supported this work. This facility is supported in part by the Neural Imaging Center as part of an NINDS P30 Core Center grant #NS072030. This work was supported in part by a Harvard Medical School Fix Fund Postdoctoral Fellowship to M.A.T and by US National Institute of Health grants to E.N.L and E.A.K (R35 GM126994-01 and R35 GM118137).

## Contributions

MAT and E.N.L conceived the study and M.A.T designed the study. M.A.T performed immunohistochemistry and imaging, collection of RNA-seq samples and analysis of RNA-seq DEG results. L.X contributed to sample collection for RNA-seq experiments and immunohistochemistry, E.N contributed in size experiments, A.M.C and JK contributed in RNA-seq data analysis. M.A.T prepared figures and wrote the original draft while M.A.T and E.N.L curated the manuscript with input from all authors.

## Supplementary figures

**Supplementary Figure 1.**
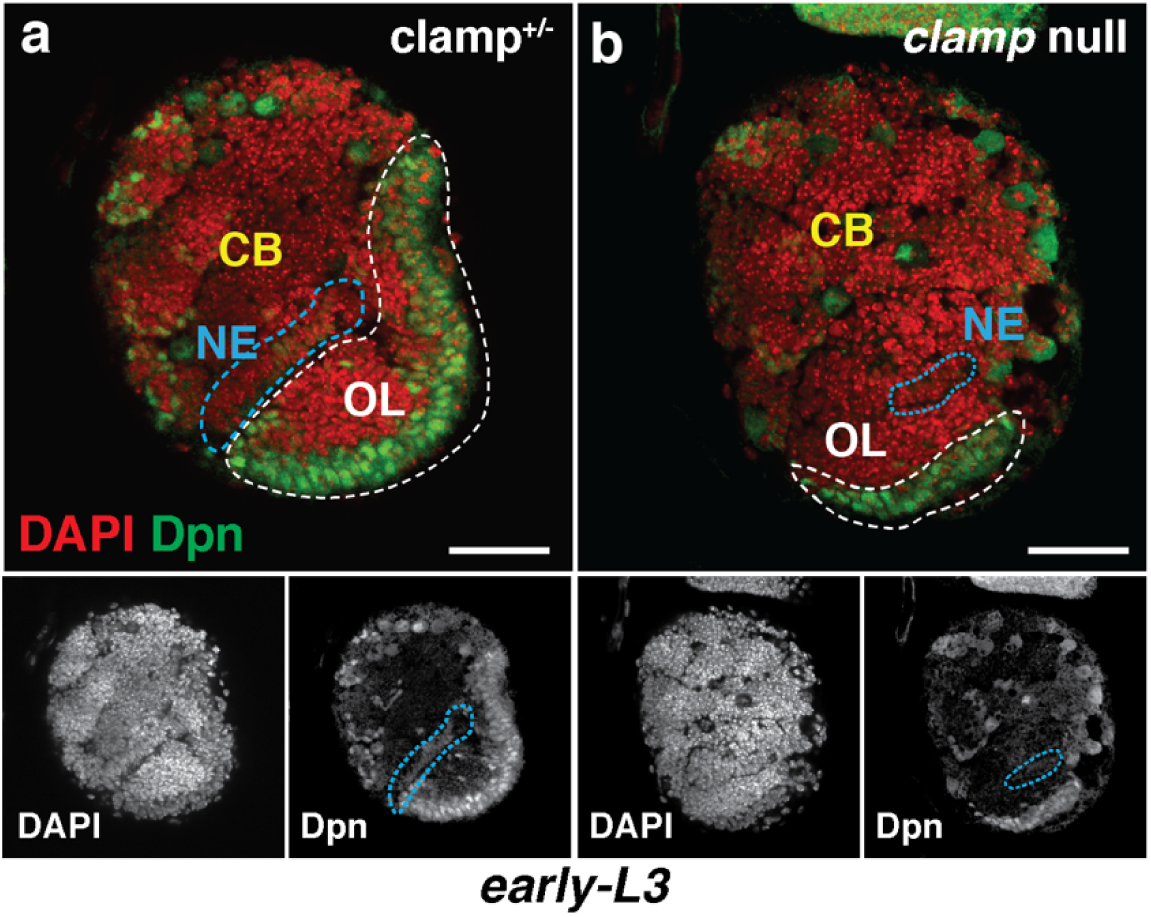
A single dose of the *Clamp* gene is sufficient to allow for normal brain development.

**Supplementary Figure 2.**
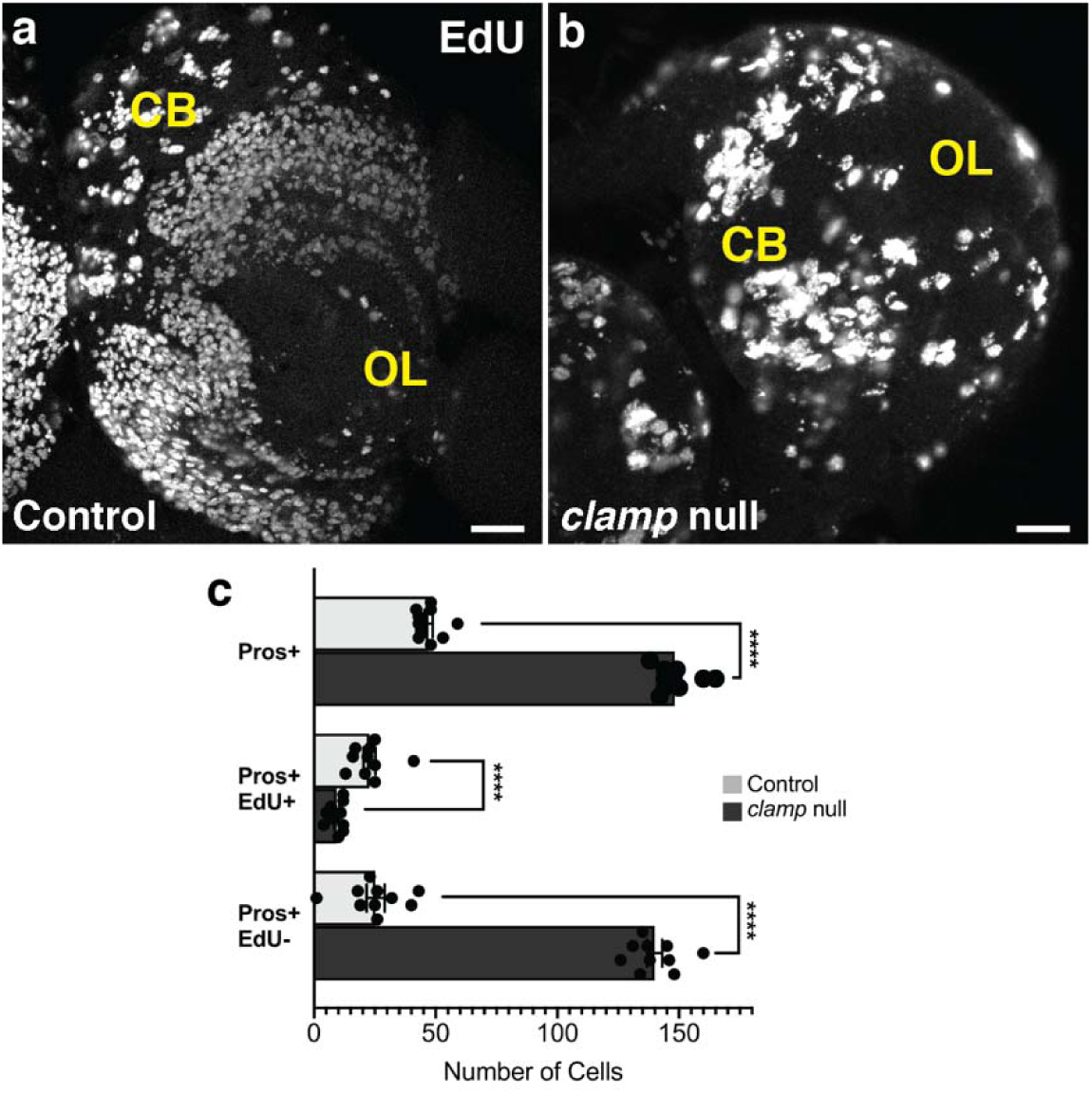

**Supplementary Figure 3.**
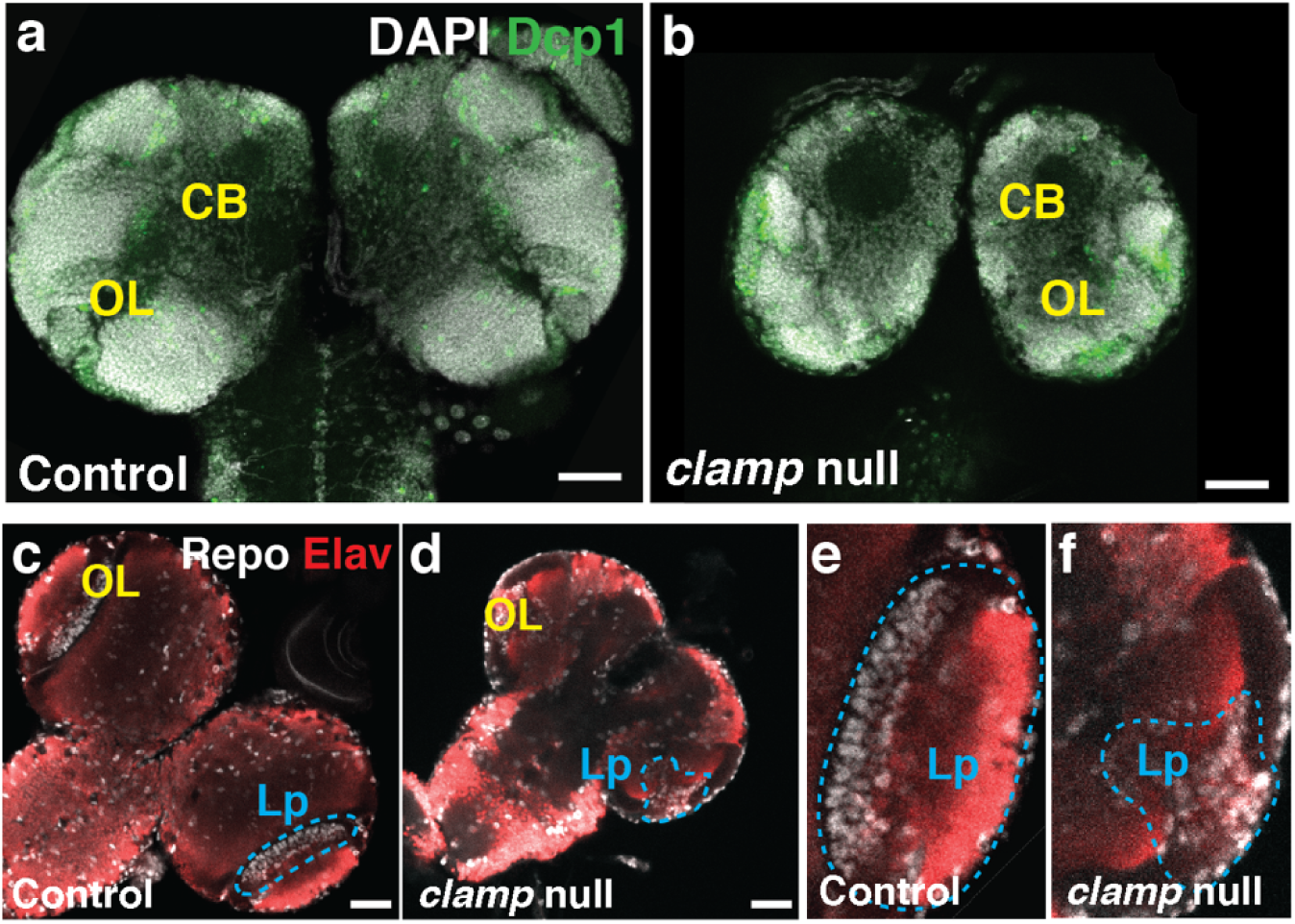

